# “Surviving and Thriving”: Evidence for Cortical GABA Stabilization in Cognitively-Intact Oldest-Old Adults

**DOI:** 10.1101/2023.09.08.556410

**Authors:** MK Britton, G Jensen, RA Edden, NA Puts, SA Nolin, SS Merritt, RF Rezaei, M Forbes, KJ Johnson, PK Bharadwaj, MK Franchetti, DA Raichlen, CJ Jessup, GA Hishaw, EJ Van Etten, AT Gudmundson, S Murali-Manohar, H Cowart, TP Trouard, DS Geldmacher, VG Wadley, N Alperin, BE Levin, T Rundek, KM Visscher, AJ Woods, GE Alexander, RA Cohen, EC Porges

**Affiliations:** Department of Epidemiology, College of Public Health and Health Professions & College of Medicine, University of Florida, Gainesville, Florida, United States; Center for Cognitive Aging and Memory, Evelyn F. and William L. McKnight Brain Institute, University of Florida, Gainesville, Florida, United States; Department of Psychology, Reed College, Portland, Oregon, United States; Russell H. Morgan Department of Radiology and Radiological Science, School of Medicine, Johns Hopkins University, Baltimore, Maryland, United States; F.M. Kirby Research Center for Functional Brain Imaging, Kennedy Krieger Institute, Baltimore, Maryland, United States; Department of Forensic and Neurodevelopmental Sciences, Institute of Psychiatry, Psychology, and Neuroscience, King’s College London, London, United Kingdom; MRC Centre for Neurodevelopmental Disorders, King’s College London, London, United Kingdom; Department of Neurology, College of Medicine, Medical University of South Carolina, Charleston, South Carolina, United States; Department of Neurology, Miller School of Medicine, University of Miami, Miami, Florida, United States; Evelyn F. McKnight Brain Institute, Miami, Florida, United States; Department of Clinical and Health Psychology, College of Public Health and Health Professions, University of Florida, Gainesville, Florida, United States; Center for Cognitive Aging and Memory, College of Public Health and Health Professions, University of Florida, Gainesville, Florida, United States; Department of Psychology, College of Science, University of Arizona, Tucson, Arizona, United States; Evelyn F. McKnight Brain Institute, Tucson, Arizona, United States; VA Boston Healthcare System, Boston, Massachusetts, United States; Department of Biological Sciences, College of Letters, Arts, and Sciences, University of Southern California, Los Angeles, California, United States; Department of Neurology, College of Medicine, University of Arizona, Tucson, Arizona, United States; Department of Psychiatry, College of Medicine, University of Arizona, Tucson, Arizona, United States; Department of Neurobiology, Heersink School of Medicine, University of Alabama at Birmingham, Birmingham, Alabama, United States; Evelyn F. McKnight Brain Institute, Birmingham, Alabama, United States; Department of Biomedical Engineering, College of Engineering, University of Arizona, Tucson, Arizona, United States; Department of Neurology, Heersink School of Medicine, University of Alabama at Birmingham, Birmingham, Alabama, United States; Heersink School of Medicine, University of Alabama at Birmingham, Birmingham, Alabama, United States; Department of Radiology, Miller School of Medicine, University of Miami, Miami, Florida, United States

## Abstract

Cortical GABA levels are reduced in older age; age-related differences in GABA may be associated with age-related cognitive change. The nature of age-related GABA differences in the highest-functioning stratum of the oldest-old (85+) population is not yet known. We extend our previously-reported Individual Participant Data Meta-Analysis of GABA levels (Porges et al., 2021) across the lifespan with four novel datasets sampling the cognitively-intact oldest-old. The slope of age-related GABA differences in cognitively-intact oldest-old adults flattens after roughly age 80. We interpret these findings as an effect of survivorship: inclusion in the study required intact cognition, and too great a reduction of GABA levels may not be compatible with neurophysiological function needed for intact cognition. This work contributes to a growing body of evidence suggesting that successful cognitive aging may require intact GABAergic function, as well as further characterizing successful aging amongst oldest-old adults.

## INTRODUCTION

Gamma-aminobutyric acid (GABA) regulates inhibitory function in the mammalian brain. Substantial evidence for reduced GABAergic function during the aging process, regarding both synthesis and receptor expression, has been reported in human and animal models (e.g., McQuail, Frazier, & Bizon, 2015; Rozycka and Liguz-Lecznar, 2017). Cross-sectional magnetic resonance spectroscopy (MRS) studies in humans have shown reduced regional GABA in older adults (Gao et al., 2013; Porges et al., 2017; Cuypers et al., 2020; Chamberlain et al., 2021; for contrast, see Pitchaimuthu et al., 2017). Recently, our group demonstrated that GABA declines gradually and nonlinearly with age (Porges et al., 2021).

Lower cortical GABA levels are associated with worse global cognition, fluid processing, sensorimotor performance, and memory performance in normative aging (Porges et al., 2017; Simmonite et al., 2019; Cassady et al., 2019; Murari et al., 2020; Jiménez-Balado et al., 2021; reviewed in Li et al., 2022). However, few studies of cortical GABA levels have included oldest-old adults (85 years and older), and none have compared young-old and old-old adults (Porges et al., 2017; Simmonite et al., 2019). The oldest-old, particularly the cognitively-intact oldest-old, may disproportionately carry longevity-related physiological traits or engage in protective behaviors (Silverman and Schmeidler et al., 2018; Kawas, Legdeur, & Corrada, 2021); “successful agers” may thereby be atypically resistant to neuropathology or able to compensate for neuropathology (e.g., Berlau et al., 2009). Given the association between GABA and cognition in normative aging, maintained GABAergic function may be protective in the cognitively-intact oldest-old (e.g., by reducing excitotoxic injury).

We describe GABA levels among cognitively-intact oldest-old adults, incorporating four novel datasets into a previously-reported individual participant data meta-analysis (Porges et al., 2021). We report stabilization in the slope of age-related GABA decline among the cognitively-intact oldest-old, potentially related to survivorship in this subpopulation.

## RESULTS

To test the linearity of GABA-age relationships in oldest-old adults, we fit separate linear regression models (Table 1); data were feature-scaled by dividing raw scores by their geometric mean to increase comparability between datasets. No model explained more than 10% of variance per R^2^, consistent with nonlinear relationships or high inter-individual variability.

**Table 1.** Regression statistics for simple linear fits.

| Site | Mean Slope | 95% Credible Interval of Posterior Estimate | Mean residual $\sigma$ | Lower and upper bounds | $R^2$ |
| --- | --- | --- | --- | --- | --- |
| Site 1 | 0.0101 | -0.0083 to 0.0289 | 0.195 | 0.1539 to 0.2510 | 0.037 |
| Site 2 | 0.0166 | -0.0148 to 0.0499 | 0.226 | 0.1578 to 0.3313 | 0.077 |
| Site 3 | -0.0125 | -0.0289 to 0.0036 | 0.1424 | 0.1083 to 0.1912 | 0.091 |
| Site 4 | 0.0143 | -0.0344 to 0.0615 | 0.1954 | 0.1398 to 0.2763 | 0.022 |

To characterize nonlinear GABA trajectory in oldest-old age, we fit a nonlinear penalized basis spline model incorporating data previously reported in Porges et al. (2021); the mean spline and credible interval are shown in Figure 1. The first derivative of the basis spline model was also generated, reflecting the change in velocity of GABA+ decline in oldest-old age; the first derivative after age 60 is inset in Figure 1.

**Figure 1.**
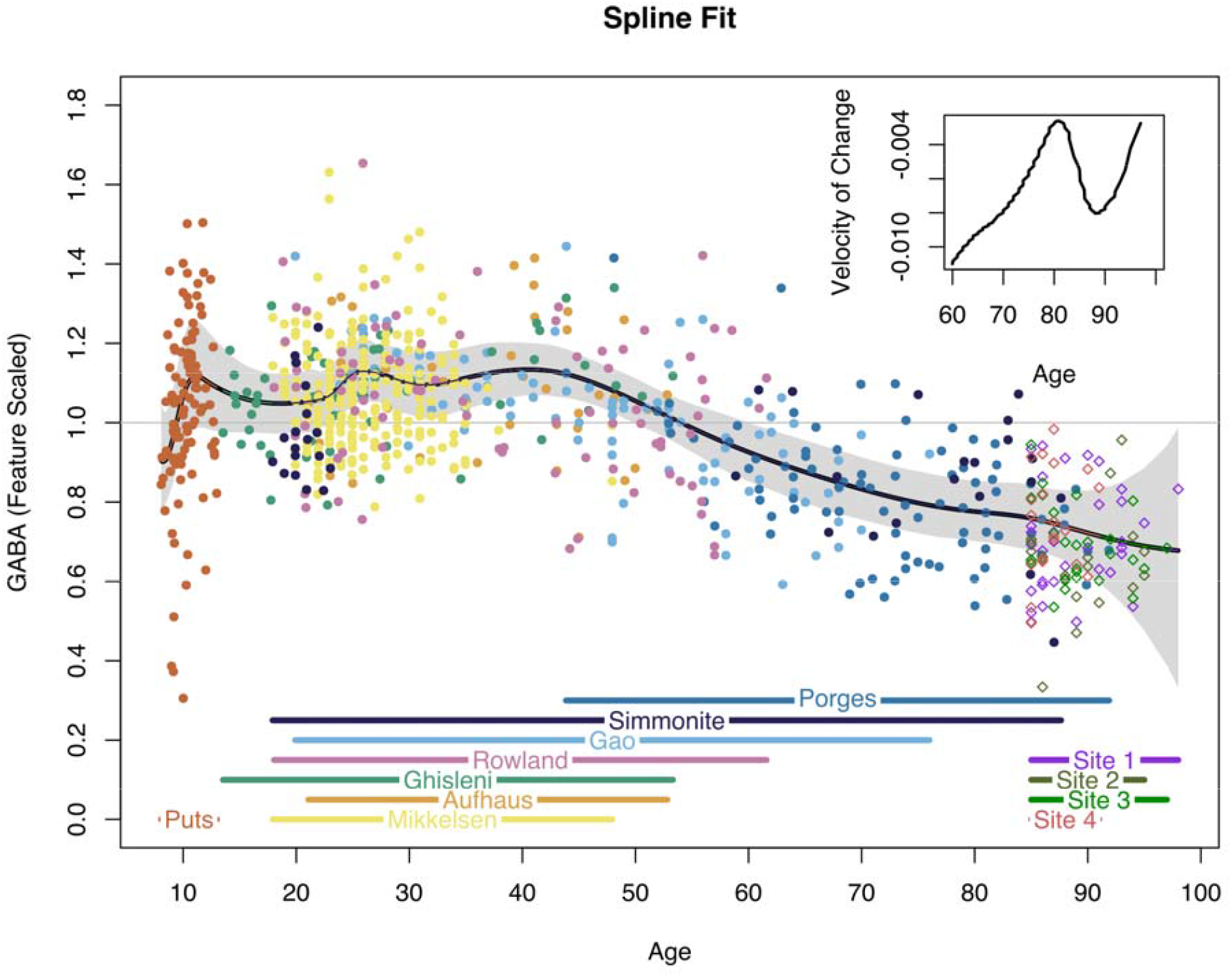
Penalized basis spline model of cortical GABA concentrations across the lifespan. The inset figure depicts a portion of the first derivative of the basis spline model, reflecting the change in steepness of the GABA slope with increasing age.

We additionally examined posterior estimates of the relative scaling factor *F*_*s;*_ *F*_*s*_ represents the dataset-specific scaling factor required to bring methodologically heterogeneous data into common units (see Methods). Estimated *F*_*s*_ for each dataset, log-scaled to emphasize ratios and centered to mean *F*_*s*_, is plotted in Figure 2. Estimates of Fs for the four Cr-referenced novel datasets, like estimates from the three other Cr-referenced datasets (Mikkelsen, Gao, and Simmonite), were smaller than estimates for water-referenced datasets, suggesting consistency between datasets using similar methodologies.

**Figure 2.**
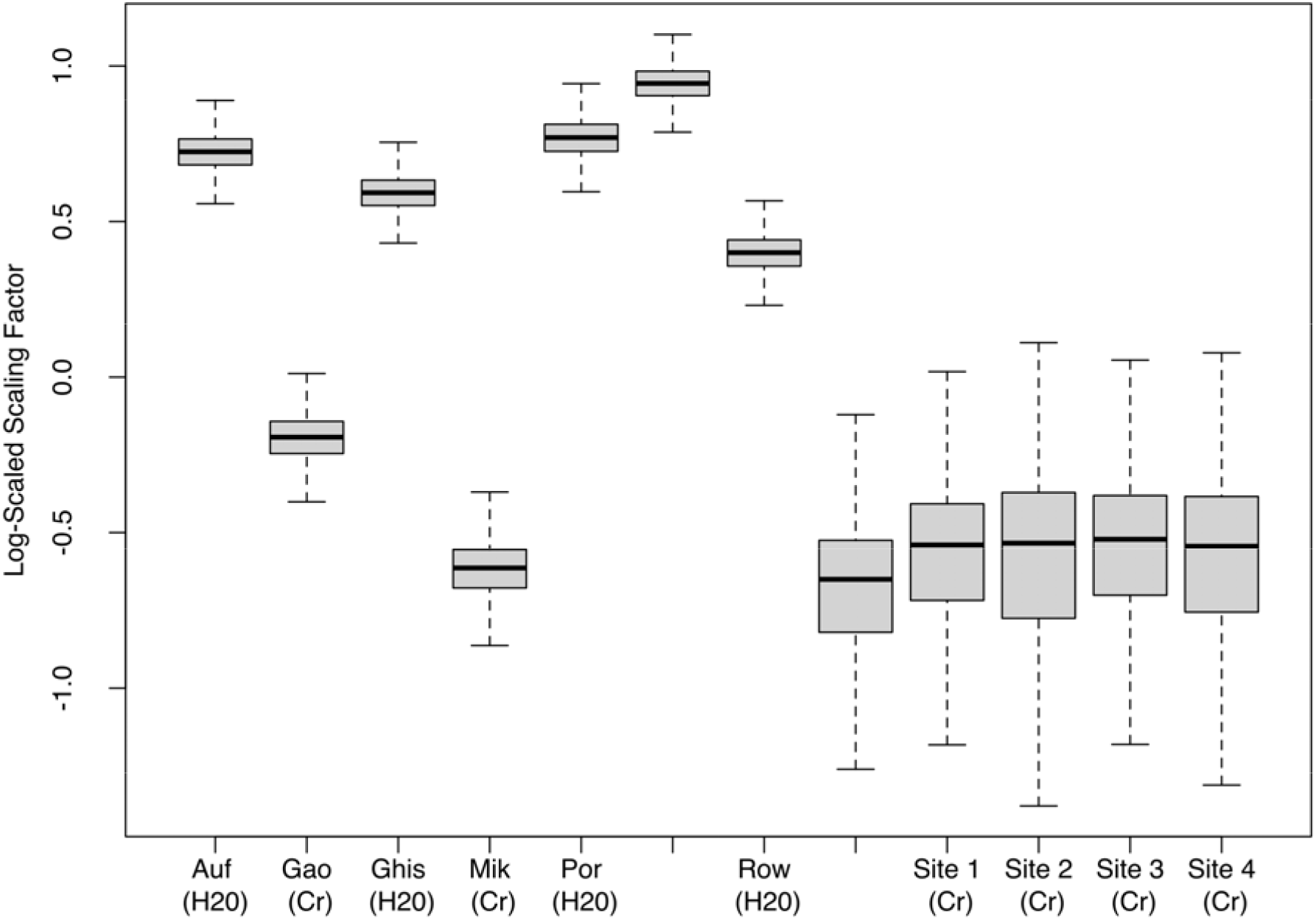
Log-scaled posterior estimates of the feature scaling factor *F*_*s*_ for each dataset assessed with the spline model.

## DISCUSSION

Although GABA declines with age and may be implicated in sustaining normal cognitive performance, few studies have examined age-related GABA differences in oldest-old adults (85+). Our individual participant data meta-analysis revealed that the slope of age-related differences in cortical GABA appears to flatten in cognitively-intact oldest-old adults; supporting this finding, the velocity of change in GABA decreases after age 80, apparently reflecting the introduction into the model of a large number of cognitively-intact oldest-old individuals in this age group, and only begins to increase again by 90. In the context of a positive GABA-cognition association, this descriptive finding is consistent with the possibility of a GABA “floor” below which cognitive function is unlikely to remain intact in oldest-old age.

The reported effect may be driven in part by survivorship and sample selection. All participants met an inclusion threshold of 22 or higher on the MoCA, reflecting grossly-intact cognitive performance. Successfully-aging oldest-old adults are a subset of a highly-selected, atypical minority of their birth cohort. Most individuals do not reach the age of 85, and those who do may be physiologically or behaviorally unusual (Silverman and Schmeidler et al., 2018; Kawas, Legdeur, & Corrada, 2021); even within oldest-old adults, an estimated 18-35% live with dementia (Gardner, Valcour, & Yaffe, 2013) and an additional 16-23% live with Mild Cognitive Impairment (Peltz et al., 2012; Yaffe et al., 2011). That is, with increasing age, the normative adult experiences accumulating age-related changes and neuropathological burden and concomitant cognitive decline, ultimately falling below a threshold required to sustain functional performance (Hertzog et al., 2008). The hypothesized relationship to GABA is visualized in Figure 3: smoothed extracted spline data are superimposed over the conceptual model proposed by Hertzog (2008). The trajectory of GABA across the lifespan shows good agreement with age-related cognitive decline, consistent with GABA levels representing a physiological constraint, below which cognition might transition to a dysfunctional state. Conversely, the attenuated slope of age-related GABA differences in our participants suggests that our participants show the range of GABA levels expected in the absence of significant age-related cognitive dysfunction. We characterize this as a “surviving and thriving” effect, in which the healthy and high-performing individuals included in a nonclinical sample exist within a relatively small range of variation of the outcome of interest.

**Figure 3.**
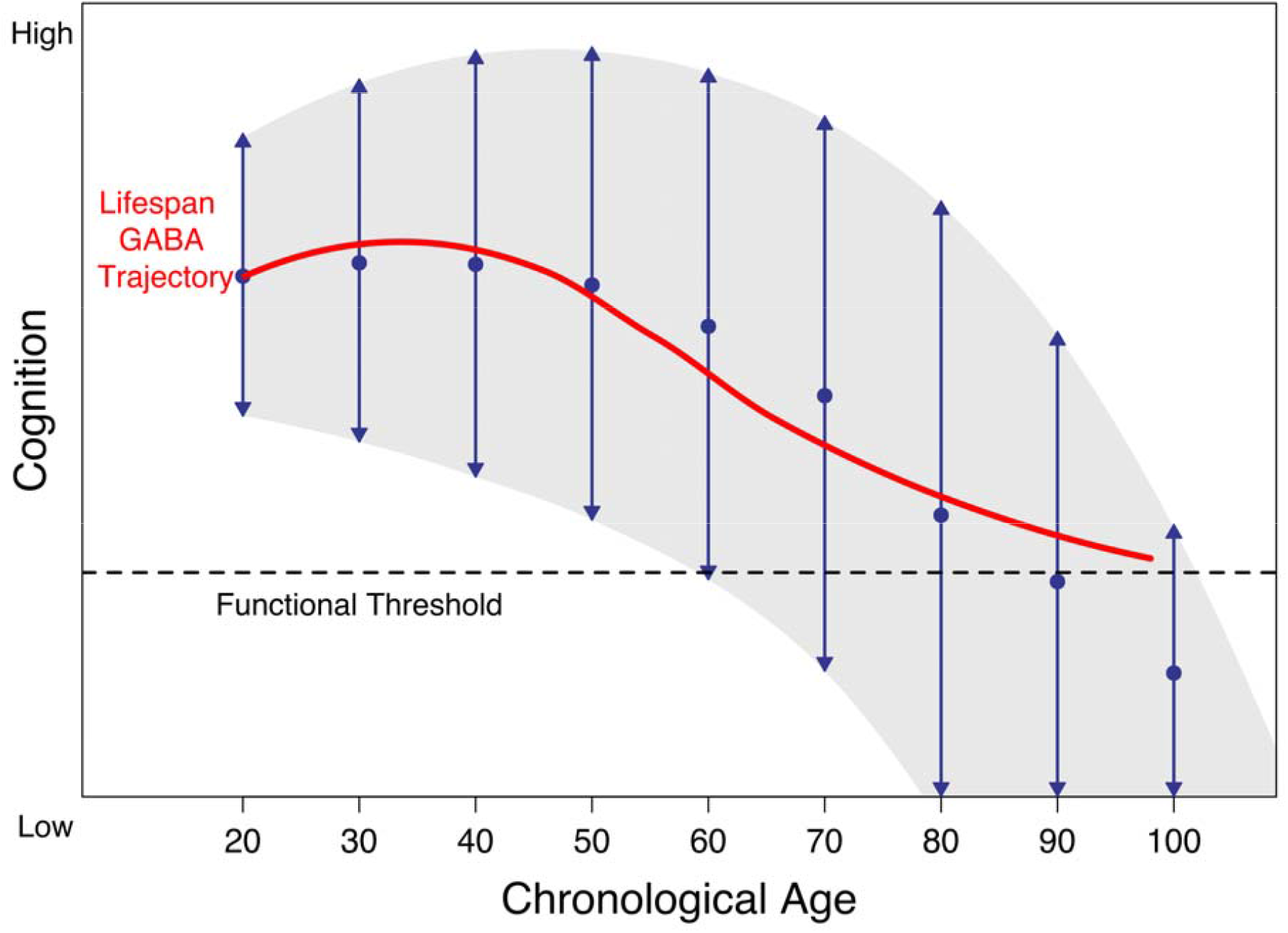
Smoothed GABA spline data are superimposed over a conceptual model of age-related cognitive change. Age-associated differences in GABA may function as a biological influence modulating trajectories of cognition in aging (Hertzog et al., 2008). Figure adapted from Hertzog et al. (2008).

While MRS does not identify the mechanism underlying age-related decline in GABA, preclinical and postmortem research have identified potential cellular and molecular drivers. GABAergic interneurons and GABA-synthesizing enzymes GAD65 and GAD67 are depleted in rodent models of aging (reviewed in Rozycka and Liguz-Lecznar, 2017), and cortical GAD65 is reduced in older adults in postmortem human studies (Pinto et al., 2010). Depletion of GABA may in turn disrupt excitatory-inhibitory balance, leaving the brain vulnerable to excitotoxic injury.

Strengths of our analysis include the individual participant data meta-analytic approach, which allows us to leverage datasets covering the full lifespan; additionally, our Markov Chain Monte Carlo approach, in which dataset-specific feature scaling factors and the spline model are estimated simultaneously, reduces the impact of inter-dataset methodological heterogeneity on our model. However, our analysis has several limitations. First, although our model represents the full lifespan, all included data are cross-sectional. Therefore, we cannot report individual participants’ cognitive trajectories; some participants with currently-intact cognitive function may have borne neuropathology undetected due to strong compensatory abilities or high baseline function (Stern, 2009) and may ultimately experience cognitive decline, while others may sustain cognitive function throughout their lives. Second, although all participants were cognitively-intact, we did not directly examine association between GABA and cognitive performance; while prior evidence from mixed young-old and oldest-old samples (e.g., Porges et al., 2017) is consistent with an association, it is also possible that, given sufficient GABA levels to maintain functioning, other aspects of brain health may explain more variance in cognition in the oldest-old. This possibility remains to be tested. Third, we did not compare our cognitively-intact sample to oldest-old adults with MCI or dementia; we hypothesize that impaired oldest-old adults would show lower GABA levels. Fourth, because individual sex, race/ethnicity, and education data were unavailable for the eight published datasets, we were unable to assess the impact of demographic characteristics on our findings. Prior studies have demonstrated that the impact of age on GABA may be greater in women (Gao et al., 2013) and that GABA may be associated with cognition in older women, but not older men (Jiménez-Balado et al., 2021). Furthermore, our sample was predominantly White and well-educated. Further research should consider sex, and potentially environmental factors such as socioeconomic status, directly. Additionally, potential lifestyle influences on GABA, such as lifetime substance use, are beyond the scope of the present analysis. Finally, we analyzed only a small number of participants over the age of 95, resulting in greater uncertainty regarding the trajectory of GABA differences after 95 (and a wider credible interval).

In conclusion, we observed attenuation of the slope of age-related differences in cortical GABA concentrations in oldest-old age, consistent with a minimum GABA “floor” physiologically necessary to sustain cognitive function. This lower limit may also reflect a state below which excitotoxic processes accelerate. The current analysis provides a descriptive benchmark for future studies examining GABA change in the oldest-old population; additionally, this analysis highlights the impact of study inclusion criteria and survivorship on observed physiological stability in oldest-old age.

## MATERIALS AND METHODS

### Design

206 community-dwelling adults aged 85 to 99 were recruited at the University of Alabama at Birmingham, the University of Florida, the University of Miami, and the University of Arizona as part of a larger study of cognition and brain function in oldest-old adults (McKnight Brain Aging Registry), funded by the Evelyn F. McKnight Brain Foundation. Sites have been designated Site 1-Site 4 to protect participant anonymity. Exclusion criteria included Mild Cognitive Impairment (MCI) or dementia, a history of neurological disease, and a history of intractable psychiatric disorders. Participants were screened for MCI by research coordinators using the Telephone Interview for Cognitive Status-Modified (TICS-M) and Montreal Cognitive Assessment (MoCA): scores of 28 or below on the TICS-M and 22 or below on the MoCA resulted in a consensus conference to assess cognitive performance, aided by a neurologist’s evaluation. All participants gave informed consent and were compensated for their participation. Study procedures were approved by the Institutional Review Boards of the University of Alabama at Birmingham (#X160113004), University of Florida (#201300162), University of Miami (#20151783) and University of Arizona (#1601318818) and conformed to the standards of the Declaration of Helsinki.

Of the 206 participants recruited, MEGA-PRESS MRS data were acquired from 164 individuals. 39 scans were corrupted or failed visual quality inspection. To ensure that all participants were cognitively-intact, an additional 24 participants scoring 22 or below on the MoCA were excluded from analysis; a cutoff score of 23 has been recommended to minimize the false positive rate for aMCI (Carson, Leach, & Murphy, 2018). An additional participant was excluded due to a GABA+/Cr value falling 4.78 standard deviations from the site mean. A GABA+/Cr fit error criterion of 15% was applied; however, this did not result in the exclusion of additional participants. Demographic data for retained participants (N = 100) are presented in Table 2.

**Table 2.** Characteristics of McKnight Brain Aging Registry participants retained in final analysis.

| Characteristic | Overall,<br>N = 100<sup>1</sup> | Site 1,<br>N = 34<sup>1</sup> | Site 2,<br>N = 18<sup>1</sup> | Site 3,<br>N = 28<sup>1</sup> | Site 4,<br>N = 20<sup>1</sup> |
| --- | --- | --- | --- | --- | --- |
| Age | 88.66 (3.35) | 88.82 (3.62) | 89.72 (3.53) | 89.07 (3.38) | 86.85 (1.87) |
| Years of Education | 16.18 (2.96) | 15.59 (2.79) | 15.89 (2.47) | 17.21 (3.14) | 16.00 (3.23) |
| Sex |  |  |  |  |  |
| Male | 41 (41%) | 17 (50%) | 8 (44%) | 8 (29%) | 8 (40%) |
| Female | 59 (59%) | 17 (50%) | 10 (56%) | 20 (71%) | 12 (60%) |
| Race |  |  |  |  |  |
| White | 95 (95%) | 32 (94%) | 18 (100%) | 28 (100%) | 17 (85%) |
| Black | 4 (4.0%) | 2 (5.9%) | 0 (0%) | 0 (0%) | 2 (10%) |
| Asian | 1 (1.0%) | 0 (0%) | 0 (0%) | 0 (0%) | 1 (5.0%) |
| MoCA Score | 25.55 (1.94) | 25.26 (1.86) | 24.83 (1.98) | 26.36 (1.97) | 25.55 (1.70) |

<sup>1</sup>Mean (SD); n (%)

### Imaging

GABA+-edited MEGA-PRESS data were acquired on Siemens 3T scanners (Skyra or Prisma) at each site, using the Siemens MEGA-PRESS WIP. Voxels were 30 × 30 × 30 mm^3^. The voxel was placed on the frontal midline, superior to the genu of the corpus callosum. A transversal 20 mm saturation band was placed along the skull, 1-2 mm away from the top of the voxel. Scans were edited for GABA + macromolecules, placing ON editing pulses at 1.9 ppm and OFF at 7.46 ppm; the TE was 68 ms and the TR 2000 ms; 320 averages were acquired (160 ON and 160 OFF). The total scan took 10:48 minutes. A total of 4096 data points were collected. The spectral width was 4000 Hz.

Data were analyzed in Gannet 3.3.1 (Edden et al., 2014), implemented in MATLAB 2022a. Cr was used as the reference signal. No deviations were made from automated procedure. To visualize voxel placement, a heatmap depicting average voxel location normalized to and superimposed over the MNI152 template brain was generated using Osprey 2.5.0 (Oeltzschner et al., 2020) (Figure 4). A representative difference spectrum is shown in Figure 5. Full details of the analytic pipeline, including GABA SNR, linewidth, and fit error(%) are reported in Supplementary Table 1 per consensus recommendations (Lin et al., 2021).

**Figure 4.**
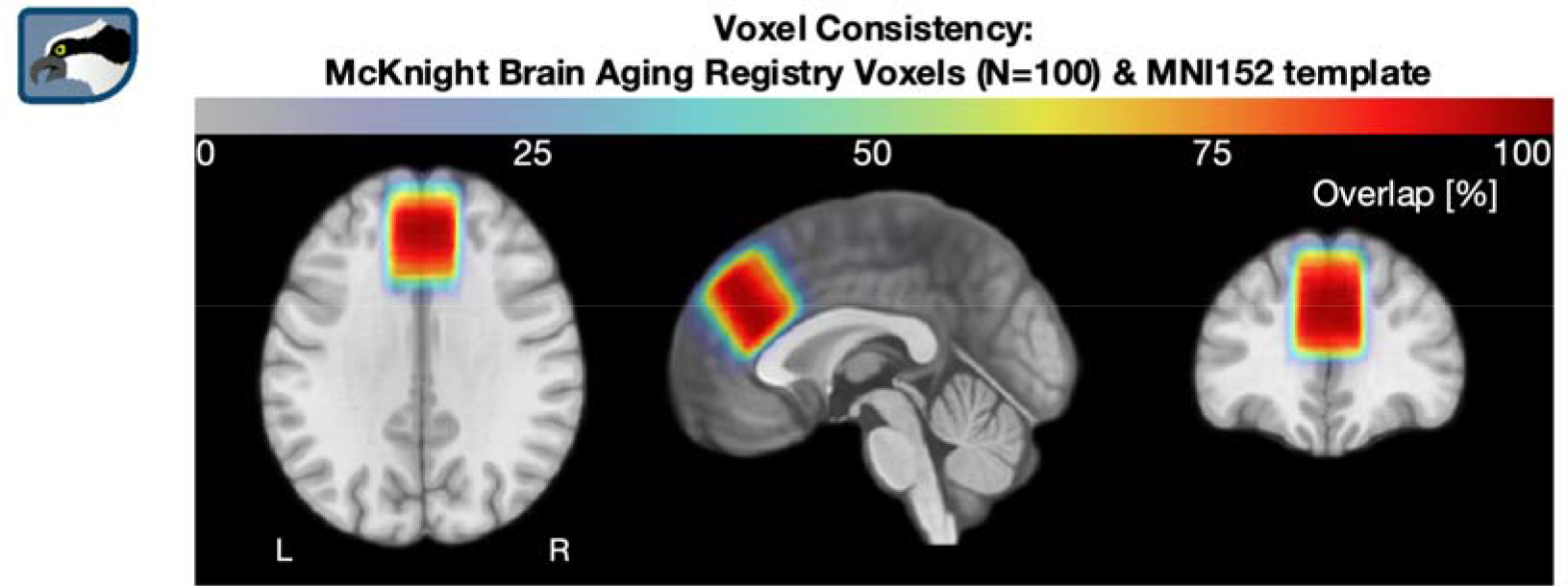
Heatmap of frontal voxel location, superior to the genu of the corpus callosum, superimposed over the MNI152 template brain.

**Figure 5.**
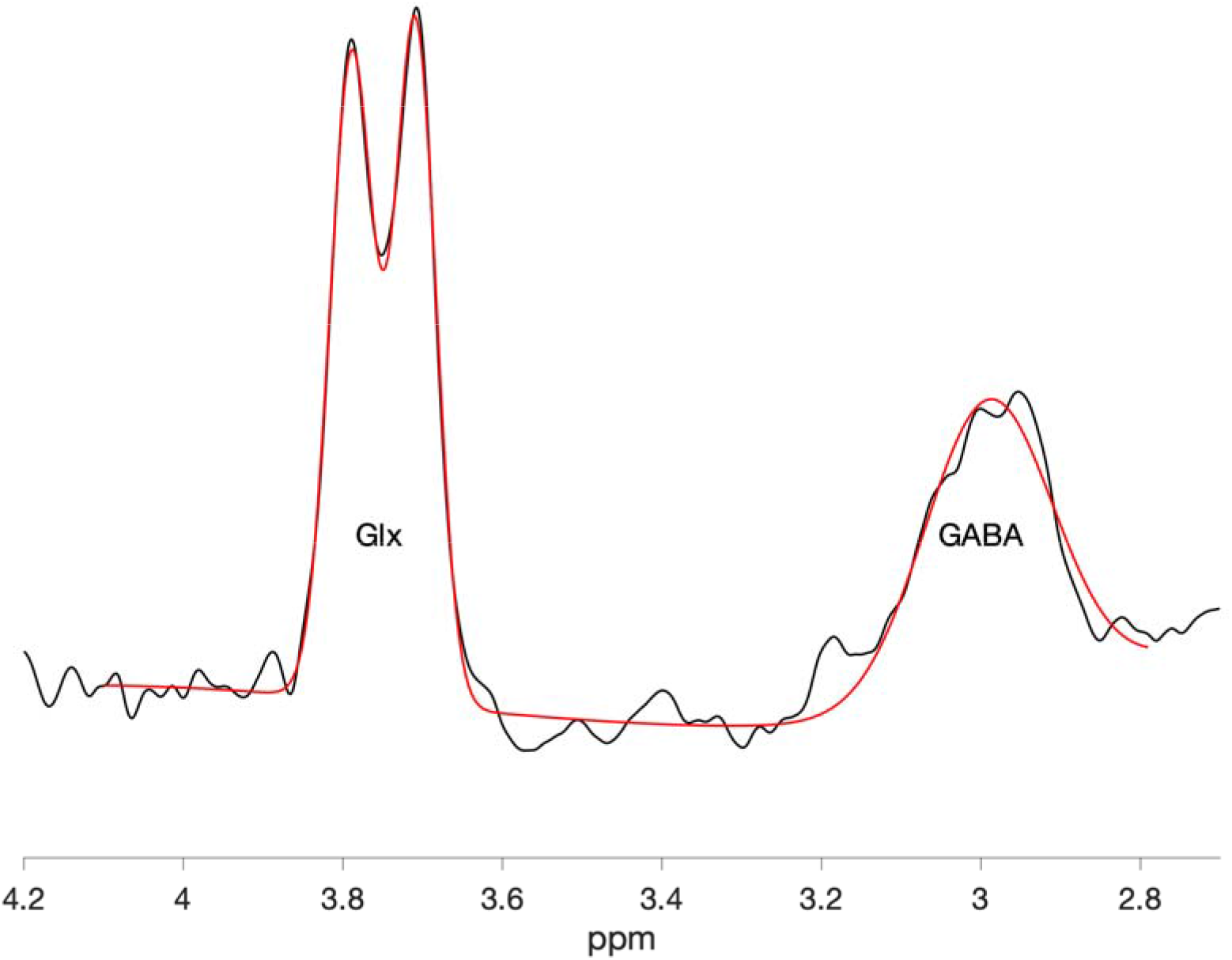
GABA+/Cr and Glx/Cr difference spectrum from a representative participant. The raw difference spectrum is plotted in black; the fitted model is plotted in red.

### Statistical Analysis

Eight other datasets were derived from a systematic literature search identifying studies reporting cortical GABA, acquired using MEGA-PRESS, in healthy populations. The methods and results of this systematic review have previously been fully reported in Porges et al. (2021).

All analyses were conducted in R 4.2.2 (R Core Team, 2022) using the cmdstanr package to implement Stan 2.21.0 (Stan Development Team, 2021). Following the approach of Porges et al. (2021), posterior probability distributions were estimated for all parameters simultaneously using Markov Chain Monte Carlo (MCMC). This approach permits estimates to covary appropriately for non-linear regression. For each model, we fit the overall function, a global error term, and one feature scaling factor *F*_*s*_ per dataset. *F*_*s*_ is the estimated multiplier needed to bring each dataset into standardized units with an overall geometric mean of 1.0, thereby correcting for systematic methodological differences. This approach assumes that data within a single dataset are comparable, that all data reflect a similar underlying lifespan trajectory, and that age ranges overlap between datasets. We generated *F*_*s*_ as in Porges et al. (2021), weakly penalizing to discourage unlikely extreme values at the tails of the distribution and excluding negative values by setting a lower bound.

We initially fit separate linear regressions in Stan predicting GABA+/Cr+PCr concentrations from age for each of the four novel datasets. We then fit a penalized cubic basis spline model (Kharratzadeh 2017; Porges et al. 2021), which imposes few assumptions on the shape of the dataset. 19 knots were spaced evenly throughout the model and a smoothing parameter was imposed to minimize overfitting (Kharratzadeh 2017). To quantify the change in model slope with the addition of the four novel datasets, we took the first derivative of the model, which represents the velocity of GABA+ change across the lifespan.

## FUNDING

The work was primarily supported by the McKnight Brain Research Foundation and the University of Florida Center for Cognitive Aging and Memory. Additional funding was received from NIH 5R015R01EB023963-06 and NIH 5R01AG054077-02. Mark Britton was funded through the NIH/NIAAA T32AA25877 and the University of Florida Center for Cognitive Aging and Memory.

## DATA AVAILABILITY STATEMENT

All code and previously-published data, including data previously reported in Porges et al. (2021), are publicly accessible from the Open Science Framework here: https://osf.io/ej6tv/. McKnight Brain Aging Registry data are available from the McKnight Brain Research Foundation upon reasonable request.

## Notes

https://osf.io/ej6tv/

### Competing Interest Statement

The authors have declared no competing interest.

