## Supplemental Table 1 for "“Surviving and Thriving”: Evidence for Cortical GABA Stabilization in Cognitively-Intact Oldest-Old Adults"

Supplementary Table 1: Summary MRSinMRS Report

Here we provide a summary following following minimum reporting standards in MRS generated in Osprey. For details, please see Lin et al. 'Minimum Reporting Standards for in vivo Magnetic Resonance Spectroscopy (MRSinMRS): Experts' consensus recommendations. NMR in Biomedicine. 2021;e4484. [doi.org/10.1002/nbm.4448](https://doi.org/10.1002/nbm.4448)

| 1. Hardware | |
| --- | --- |
| a. Field strength [T] | 3T |
| b. Manufacturer | Siemens |
| c. Model (software version if available) | Skyra (two sites); Prisma (two sites) |
| d. RF coils: nuclei (transmit/receive), number of channels, type, body part | 1H 64-channel head coil |
| e. Additional hardware | - |

| 2. Acquisition | |
| --- | --- |
| a. Pulse sequence | MEGA-PRESS (Mescher et al., 1998) |
| b. Volume of interest (VOI) locations | Frontal cortex, superior to genu of corpus callosum and aligned with corpus callosum. See Figure 4 for heatmap demonstrating VOI overlap across all included participants. |
| c. Nominal VOI size [mm^3^] | 30 x 30 x 30 mm^3^ |
| d. Repetition time (TR), echo time (TE) [ms] | TR 2000 ms, TE 68 ms |
| e. Total number of averages per spectrum  i. Number of averaged spectra per subspectrum | 320 total averages; 160 averages per subspectrum (ON and OFF) |
| f. Additional sequence parameters  i. Editing pulse frequencies | F1: 4000 Hz, 4096 points ppm_ON_ = 1.90, ppm_OFF_ = 7.50 |
| g. Water suppression method | - |
| h. Shimming method, reference peak, and threshold of acceptance of shim chosen | - |
| i. Trigger or motion correction | - |

| 3. Data analysis methods and outputs | |
| --- | --- |
| a. Analysis software | Gannet 3.3.1 |
| b. Processing steps deviating from Gannet | None |
| c. Output measure | GABA+/Cr |
| d. Quantification references and assumptions, fitting model assumptions | Fitting method: Nonlinear least-squares fit to GABAGlx using Gaussian peak w/ linear baseline (Gannet default) |

| 4. Data quality | |
| --- | --- |
| a. SNR (GABA), linewidth (GABA) [Hz] | Site \| SNR \| FWHM; Site 1 \| 16.14(3.79) \| 21.33(2.36); Site 2 \| 14.83(4.41) \| 20.20(3.68); Site 3 \| 17.33(3.75) \| 21.23(2.08); Site 4 \| 18.53(5.38) \| 21.25(2.66) |
| b. Data exclusion criteria | Failure of visual inspection GABA+/Cr value > 3 SDs from site mean GABA fit error(%) > 15 MoCA < 23 |
| c. Quality measures of post-processing model fitting | Site \| GABA Fit Error; Site 1 \| 6.20(1.66); Site 2 \| 7.12(2.98); Site 3 \| 6.30(2.57); Site 4 \| 5.80(1.06) |
| d. Sample spectrum | Figure 5 |
